# The *Chlamydia trachomatis* type III effector TarP coordinates a functional collaboration between the actin nucleators Formin 1 and Arp2/3 during invasion

**DOI:** 10.1101/2021.03.18.436027

**Authors:** Matthew D. Romero, Rey A. Carabeo

**Affiliations:** Department of Pathology and Microbiology, College of Medicine, University of Nebraska Medical Center, Omaha, NE

## Abstract

The obligate intracellular pathogen *Chlamydia trachomatis* manipulates the host actin cytoskeleton to assemble actin-rich structures that drive pathogen entry. This actin remodeling event exhibits relatively rapid dynamics that, through quantitative live-cell imaging, was revealed to consist of three phases – a fast recruitment phase which abruptly transitions to a fast turnover phase before resolving into a slow turnover of actin that indicates the end of actin remodeling. Here, we investigate *Chlamydia* invasion in the context of actin dynamics. Efficient invasion is associated with robust actin remodeling kinetics that results from a collaborative functional interaction between two different classes of actin nucleators – formins, including formin 1 and the diaphanous-related formins mDia1 and mDia2, and the Arp2/3 complex. Recruitment of these nucleators requires the presence of the chlamydial type III effector TarP, which enables the respective nucleating activities of formin and Arp2/3 to collaboratively generate a robust actin network. A collaborative model is supported by the observation that co-inhibition of Fmm1 and Arp2/3 further reduced both actin dynamics and invasion efficiency than either treatment alone. Furthermore, inhibition of recruitment of Fmn1 and/or Arp2/3 by deleting TarP was sufficient to similarly attenuated actin kinetics and invasion efficiency, supporting a model wherein TarP is the major contributor to robust actin remodeling via its recruitment of the two classes of actin nucleators. At the population level, the kinetics of recruitment and turnover of actin and its nucleators were linked. However, a more detailed analysis of the data at the level of individual elementary bodies showed significant variation and a lack of correlation between the kinetics of recruitment and turnover, suggesting that accessory factors variably modify actin kinetics at individual entry sites. In summary, efficient chlamydial invasion requires a specific profile of actin dynamics which are coordinated by TarP-dependent recruitment of two classes of actin nucleators.

**Author Summary:** The obligate intracellular pathogen *Chlamydia trachomatis* relies upon manipulation of the host actin cytoskeleton to drive its entry into host cells, such that impairment of actin dynamics attenuates *Chlamydia* invasion. Collaboration between two classes of actin nucleators, formin and Arp2/3, are known to enhance actin recruitment and turnover; we found that recruitment of both proteins to the signaling complex established by the type III secreted effector, TarP, was important for pathogen internalization. Furthermore, Formin 1 and Arp2/3 are co-recruited to sites of entry, and pharmacological inhibition of either actin nucleator impaired recruitment of the other, indicating a functional cooperation between branched and filamentous actin nucleation within pathogen entry sites. Disruption of this cooperation negatively impacted both actin dynamics and *Chlamydia* internalization, indicating that TarP-dependent entry of *Chlamydia* into non-phagocytic cells operates through the recruitment and activation of Arp2/3 and Formin 1. Finally, kinetic analysis of actin recruitment and turnover revealed that these processes were independently regulated, in addition to implicating the presence of local factors that fine-tune actin dynamics and subsequent invasion.

## Introduction

Infections caused by the obligate intracellular bacterium *Chlamydia trachomatis* are both the leading cause of preventable blindness and the most prevalent bacterial form of sexually transmitted disease worldwide (1). *Chlamydiae* feature a biphasic developmental cycle divided between metabolically inactive elementary bodies, which infect host cells, and metabolically active reticulate bodies, which replicate in a membrane-bound vacuole known as an inclusion (2). Invading elementary bodies initially form a reversible electrostatic attachment onto heparin sulfate proteoglycans on the host cell surface (3,4). Shortly after, *Chlamydia* engages and activates a multitude of host cell receptors while simultaneously delivering an array of bacterial effectors by a type III secretion system (5–7). Activation of host receptors and delivery of bacterial effectors during invasion allow *Chlamydia* to exploit regulatory components of the actin cytoskeleton like Arp2/3, Rho GTPases, and vinculin amongst others (5,7–11). By manipulating cytoskeletal regulators, *Chlamydia* induces the formation of actin-rich microstructures on the cell surface that facilitate engulfment of the pathogen (12,13). These structures can adopt a variety of configurations, including phagocytic caps, membrane ruffles, filopodia, and hypertrophic microvilli (9,14). A distinguishing characteristic of structures generated by *Chlamydia* during invasion is their rapid and highly localized assembly (12). Whereas *S. typhimurium* and *R. parkeri* recruit actin in a diffuse pattern over the span of 5-15 minutes (15–17), actin recruitment and localized assembly of invasion-associated structures by *Chlamydia* is completed within two minutes (18).

Each structure is assembled entirely or in part by elongated bundles of filamentous actin, which are often associated with the activity of a class of actin nucleators known as formins (19). Broadly, formins can be divided into two subclasses: diaphanous-related formins (DRFs) which must be activated by Rho-GTPases before they can serve as actin nucleators, and non-DRF formins, which do not require activation to participate in actin filament elongation (20). Once activated, formins dimerize and incorporate monomeric actin onto the barbed end of an actin filament. Formins have been demonstrated to participate in the invasion of bacterial pathogens including *L. monocytogenes*, *S. typhimurium*, and *B. burgdorferi* (21–23). Likewise, invading *C. trachomatis* is known to interact with hypertrophic microvillar structures that are enriched with filamentous actin (12), raising the possibility that *Chlamydia* also utilizes formins during invasion.

Arp2/3 is a well-known nucleator of actin filaments that is recruited during chlamydial invasion following the activation of WAVE2 and the Rho-GTPase Rac1 by the *C. trachomatis* effector TarP (9,24). siRNA depletion of WAVE2, a nucleation promoting factor that activates the Arp2/3 complex, prevented actin recruitment at *Chlamydia* entry sites and strongly reduced the efficiency of invasion, highlighting the importance of Arp2/3-mediated actin remodeling in *Chlamydia* pathogenesis. Furthermore, the actin branching activity of Arp2/3 is known to synergize with the actin elongating properties of formins, generating a dense network of branched and elongated actin filaments (25). This collaborative interaction is essential for many cellular processes like cell motility, division, and phagocytosis (26,27), and also enhances the actin remodeling capabilities of invading pathogens. For instance, the spiraling filamentous protrusions formed by *B. burgdorferi* or the raised actin-rich pedestals of enteropathogenic *E. coli* are both formed by collaboration between Arp2/3 and formin (28,29). These pathogen-associated structures are reminiscent of the structures generated during *C. trachomatis* invasion, introducing the possibility that formin and Arp2/3 also act collaboratively during invasion to engulf the pathogen.

In addition to the multitude of proteins which regulate the polymerization of the actin cytoskeleton, a handful of other proteins coordinate the depolymerization of actin (30). Typically, actin depolymerization factors target older actin filaments to generate monomeric actin, which in turn is utilized to generate new actin filaments (31). Despite their role in maintaining actin homeostasis, the activities of host actin depolymerization factors are often counteracted by bacterial pathogens (32). For instance, the *C. pneumoniae* TarP ortholog CPN0572 displaces the actin depolymerization factor ADF/cofilin to stabilize actin filaments during its invasion (33). However, it is evident that actin turnover is necessary for *Chlamydia* internalization, as treatment with the actin stabilizing compound jasplakinolide prevents invasion (34). Furthermore, actin recruitment observed in previous studies is highly transient (18), indicating that *Chlamydia* employs a mechanism that induces a shift from the recruitment and polymerization of actin toward an equally rapid depolymerization, which removes the dense network of filaments generated by *Chlamydia* during invasion. The details regarding how *Chlamydia* accomplishes actin depolymerization, or the signals that initiate this process remain unknown.

Given the importance of actin recruitment to chlamydial invasion, we investigated invasion in the context of actin remodeling kinetics. Efficient invasion is associated with distinct actin dynamics consisting of three phases: rapid recruitment, a brief fast turnover phase, and a lingering slow turnover phase. We observed that the kinetic profile of actin was largely determined by the collaborative actions of nucleators of branched and filamentous actin, and their inhibition significantly altered the recruitment and turnover kinetics of actin at invasion sites. We demonstrate that *Chlamydia* recruits several species of formin to assist with actin remodeling during invasion, consistent with a previously published RNA interference screen in Drosophila S2 cells that, in addition to components of the Arp2/3 complex, identified formin and Diaphanous being required for *Chlamydia* infection (35). We found that the activities of Arp2/3 and formins (e.g. Fmn1, mDia1/2), which respectively nucleate branched and filamentous actin, were necessary for efficient internalization of *Chlamydia* and optimal recruitment of actin at the entry site. Both Arp2/3 and Fmn1 were simultaneously recruited to the site of invasion, where they collaboratively enhanced the recruitment and turnover of actin and supported efficient invasion. Furthermore, genetic deletion of TarP in *C. trachomatis* restricted the recruitment of actin, Fmn1, and Arp2/3, resulting in a substantial impairment in pathogen internalization. As such, we conclude that TarP, Arp2/3 and Fmn1 comprise a potent actin remodeling scaffold that facilitates rapid assembly and indirectly, the disassembly of the actin network. Moreover, the actin network nucleated by Fmn1 and Arp2/3 turned over more efficiently during infection than actin networks generated by either nucleator alone. At the level of individual invading elementary bodies, we observed that rapid actin disassembly was not due to the amount of actin recruited nor dictated by the rate of actin recruitment, indicating the independent regulation of these two processes that potentially involve different combinations of signaling modules that fine tune the TarP-Arp2/3-Fmn1 function. Together, these observations provide a molecular basis for efficient invasion by *Chlamydia trachomatis*.

## Results

### Inhibition of filamentous and branched nucleation of actin at the sites of entry reduces invasion efficiency and restricts actin recruitment

*Chlamydia* drives its engulfment into non-phagocytic epithelial cells by reconfiguring host membranes into invasion-associated structures such as hypertrophic microvilli (12). These structures are associated with the actin elongating properties of formins, indicating that formins might contribute to invasion. To test this, we transfected Cos7 cells with GFP-fusion constructs of Formin 1 (GFP-Fmn1), mDia1 (mEmerald- mDia1), or mDia2 (mEmerald-mDia2) and tracked their recruitment around invading Chlamydiae (Fig. 1A). We observed that all three formin species accumulated at *Chlamydia* entry sites, indicating that they are recruited during invasion. Given that multiple formin species are recruited by *Chlamydia*, we utilized the pan-formin inhibitor SMIFH2 to determine the general role of formins during invasion. SMIFH2 is ideally suited for this purpose, since it inhibits all three formin species by blockade of their conserved formin-homology 2 (FH2) domains, thus preventing formin-mediated actin polymerization (36). Additionally, we employed the Arp2/3 inhibitor CK666, which prevents Arp2/3 branching activity by occluding the actin binding cleft between subunits Arp2 and Arp3 (37). The Arp2/3 complex is known to be required for efficient invasion (9), and thus serves as a basis for comparison regarding the potential effects of formin inhibition.

**Figure 1:**
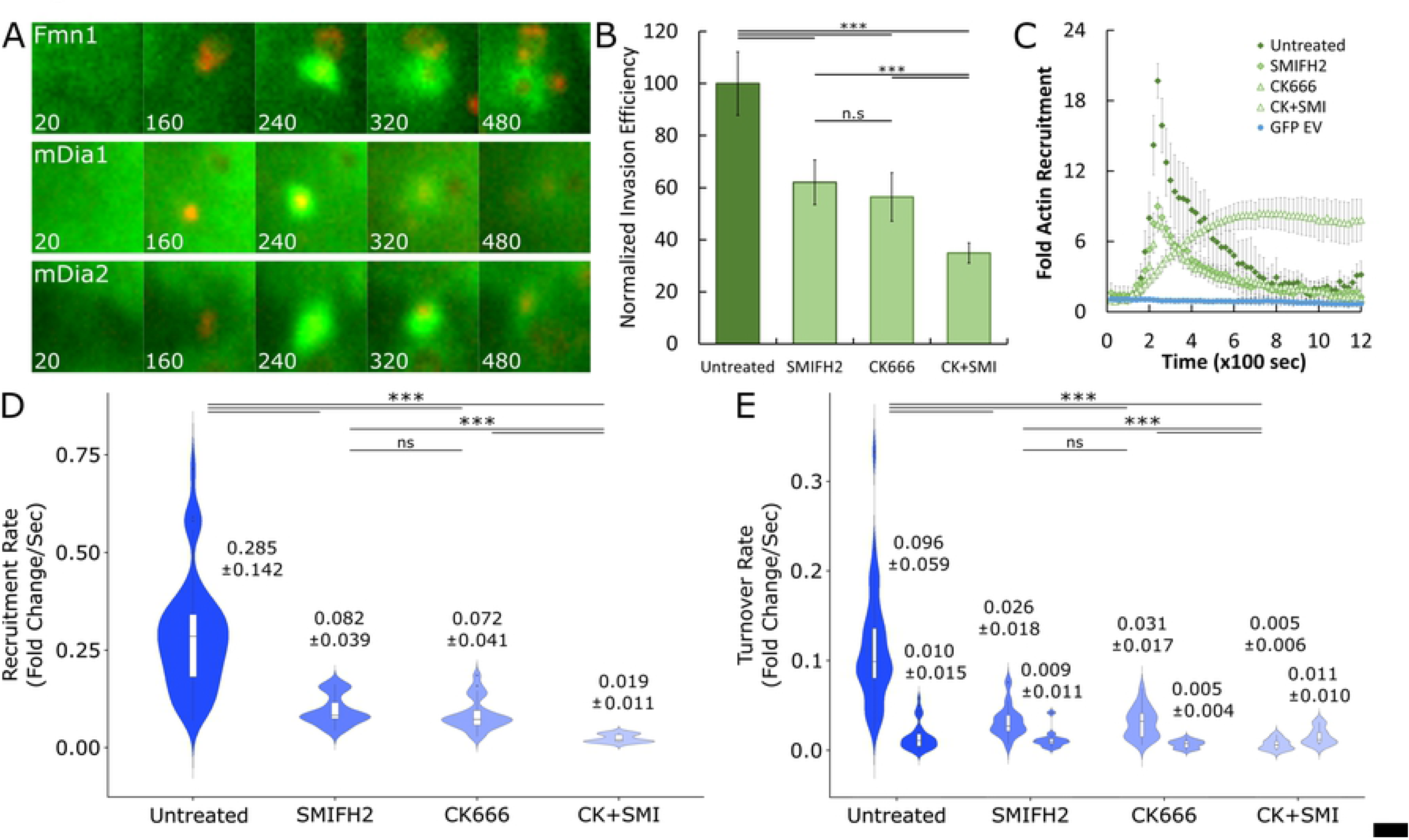
Formin and Arp2/3 are necessary for efficient invasion and optimal actin recruitment kinetics. (A) Cos7 cells were transfected with GFP-Fmn1, mEmerald-mDia1, or mEmerald-mDia2 for 24hrs prior to infection with *Chlamydia* stained with CMTPX at MOI=20. Infection was monitored by live-cell confocal microscopy using a Leica SD6000 AF Spinning Disc microscope with a 60x (NA 1.40) objective. Images were obtained every 20 seconds for 30 minutes and compiled into videos in ImageJ to identify sites exhibiting colocalization of host formins (GFP) and *Chlamydia* (RFP). (B) Cos7 cells were mock-treated or pretreated with inhibitors against formin (10µM SMIFH2), Arp2/3 (100µM CK666) or both (CK+SMI) for 1hr, infected with *C. trachomatis* serovar L2 at MOI=50 and stained using the “in-and-out” method which distinguishes non-internalized EBs from total cell-associated EBs, as described in Materials and Methods. Results were normalized against mean invasion efficiency of mock treated cells (Untreated mean = 49% invasion efficiency) and plotted as normalized mean +/- SEM. Data was collected from 20 fields, with each field containing an average of 108 Chlamydiae. Statistical significance was determined by T- test. (C) Cos7 cells were transfected with GFP-actin or a GFP empty vector plasmid for 24hrs prior to mock-treatment or pretreatment with 10µM SMIFH2, 100µM CK666, or both for 1hr. Transfected cells were infected with *Chlamydia* stained with CMTPX at MOI=20 and imaged by quantitative live-cell imaging, collecting images every 20 seconds for 30 minutes. Recruitment events were isolated by selecting regions containing CMTPX *Chlamydia* and elevated actin-GFP fluorescence. Background fluorescence was subtracted, and fold recruitment was calculated as a function of the fold increase in mean fluorescence intensity of recruited GFP-actin compared to basal GFP-actin fluorescence. Detailed visualization of this process can be found in Fig. S1. Data is displayed as mean fold recruitment for each timepoint +/- SEM compiled from a minimum N=16 recruitment events. (D,E) All actin recruitment events used to create the averaged plot shown previously (Fig. 1C) were individually divided into recruitment, fast turnover and slow turnover phases (Fig. S2). (D) Individual rates of recruitment were plotted on a violin plot with inset boxplot, reporting the median rate +/- SD for each inhibitor treatment condition. (E) Individual rates of fast turnover (Left) and slow turnover (Right) were plotted using the same method as recruitment, with fast and slow turnover plots grouped together according to condition. Violin plots contain a minimum N=16 individual rates. Statistical significance was determined by Kolmogorov-Smirnov test. All data are representative of at least 3 independent experiments, *** P ≤ 0.001.

First, we investigated whether the activities of formin and Arp2/3 are necessary for invasion by comparing the efficiency of *Chlamydia* invasion in the presence and absence of SMIFH2, CK666, or both (Fig. 1B). Cos7 cells were pre-treated with inhibitors for 60 min prior to inoculation with *C. trachomatis* serovar L2 EBs. Invasion efficiency was evaluated using the previously described “in-and-out” assay that distinguishes non-internalized EBs from total EBs associated with the infected cells (12). We normalized our data to the efficiency of invasion in absence of inhibitor in order to directly compare the effects of formin and/or Arp2/3 inhibition on invasion. Inhibiting the activity of either formin or Arp2/3 yielded a comparable impediment to invasion, each reducing invasion efficiency by roughly 40%, while simultaneous inhibition of formin and Arp2/3 reduced invasion efficiency by 60%. Together, these data suggest that while formin and Arp2/3 activity are independently important for invasion, their combined inactivation causes a more substantial impediment to pathogen entry.

To determine if this impairment in *Chlamydia* internalization is due to a deficiency in actin polymerization, we quantified the recruitment of GFP-actin at the site of invasion in the presence or absence of these inhibitors by time-lapse imaging of live Cos7 cells (Fig. 1C). Comparisons of actin recruitment between conditions were derived by calculating their respective fold recruitment values, which measures the fold increase of fluorescence intensity relative to background. A detailed account of this process can be found in the supplementary data (Fig. S1). *Chlamydia* robustly recruited actin in absence of inhibitor at an intensity of nearly 20-fold above background, which was reduced by over 10- fold following application of SMIFH2 or CK666. Interestingly, dual inhibition of formin and Arp2/3 did not further reduce actin recruitment but yielded a form of actin recruitment that was substantially slower and much more persistent, often failing to turn over throughout the duration of the experiment. Altogether, these data indicate that the activities of formin and Arp2/3 are manipulated by *Chlamydia* to enhance actin remodeling, and that inhibition of either nucleator yields a deficiency in pathogen entry.

### Recruitment and turnover of actin are slowed by inhibition of filamentous and branched actin nucleators

Our observation that inhibition of formin and Arp2/3 reduced both actin recruitment and invasion efficiency indicate a link between robust actin recruitment and pathogen internalization. Furthermore, the lack of actin turnover upon co-inhibition of formin and Arp2/3 (Fig. 1C), coupled with a drastic impairment in *Chlamydia* invasion (Fig. 1B), indicate that actin turnover is also associated with efficient internalization. Given the apparent importance of actin kinetics on the efficiency of *Chlamydia* invasion, we sought to determine how inhibition of formin and Arp2/3 alters the profile of actin recruitment. We observed that the recruitment profile of actin exhibited three distinct phases: i) recruitment, in which actin rapidly accumulates, ii) fast turnover, in which actin is rapidly lost from the sites of invasion, likely through depolymerization and iii) slow turnover, in which actin is gradually depolymerized over an extended time period (Fig. 1C). The entry and exit points for each phase were determined by plotting the first derivative values corresponding to the velocity of recruitment and turnover for each timepoint and is discussed in greater detail in the supplemental data (Fig. S2).

To compare the kinetics of actin recruitment and turnover between conditions, we employed linear regression to calculate the slope, representing the rate of recruitment and turnover for each of the three phases mentioned above, and mapped the values onto a violin plot to ascertain the distribution of rates for each inhibitor treatment (Fig. 1D, E). Inhibitor treatment reduced the rate of recruitment by 3-4 fold compared to mock-treatment (Mock=0.285 fold/sec, SMIFH2=0.082 fold/sec, CK666=0.072 fold/sec) (Fig. 1D) and altered the recruitment profile across multiple parameters, ultimately resulting in a narrower range of recruitment rates (Table 1). Together, this indicates that when only one nucleator is operational, the rate of actin recruitment is slower and more consistent. Furthermore, co-inhibition of formin and actin yielded an even more narrow range of actin recruitment rates (Range CK+SMI=0.044) and reduced the overall rate by 10-fold (CK+SMI=0.015 fold/sec). Since actin recruitment was not completely ablated by dual-inhibition of formin and Arp2/3, these data imply the presence of a formin- and Arp2/3-independent mechanism of actin recruitment at the site of invasion. However, the resultant actin network polymerizes very slowly and more uniformly than the actin network created by host actin nucleators. Collectively, this suggests that formin and Arp2/3 activity contributes to efficient invasion by enhancing the recruitment kinetics of actin, resulting in the rapid establishment of a robust actin network at the site of invasion.

**Table 1.**
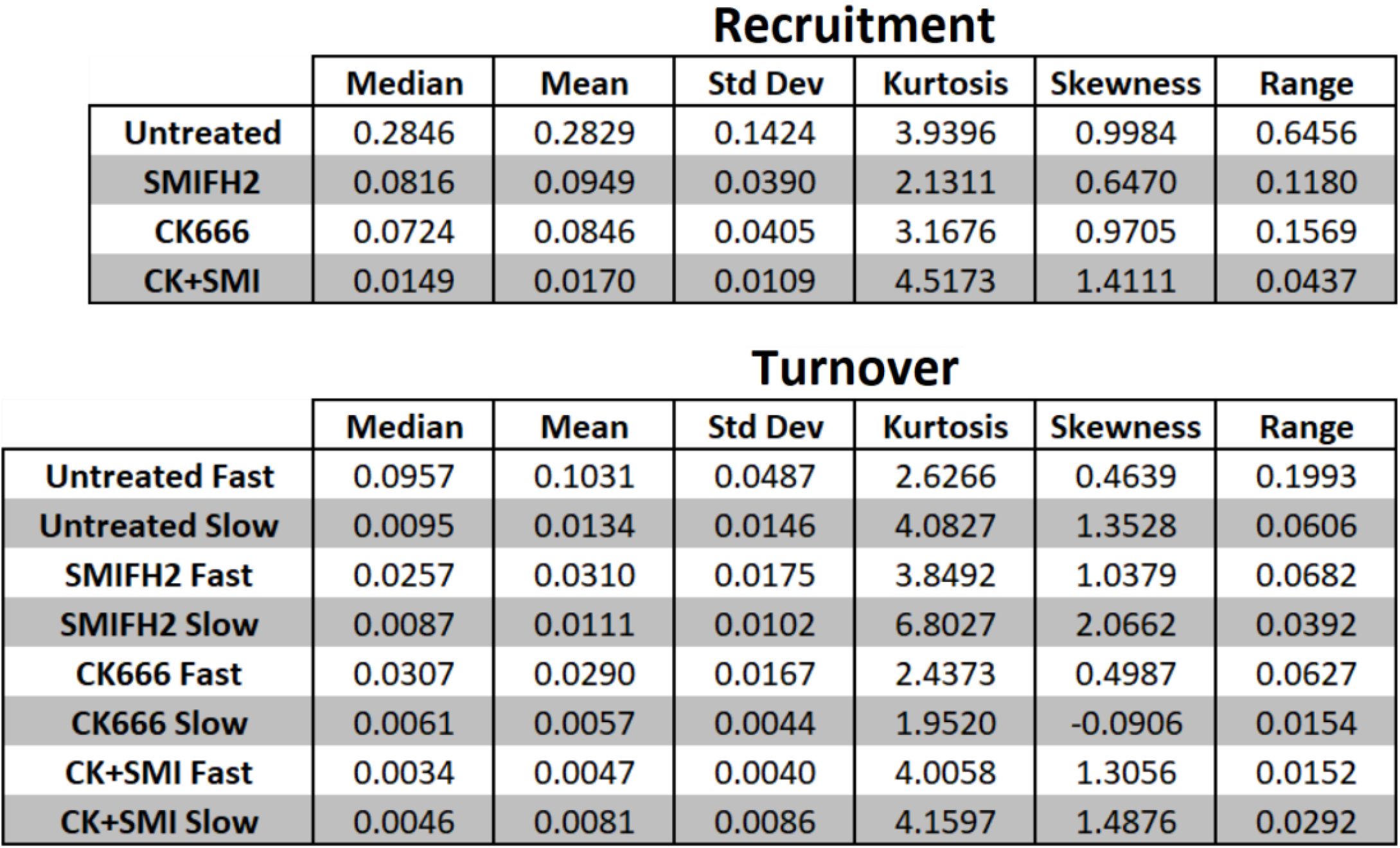
Statistical values associated with actin recruitment and turnover in presence and absence of inhibitors.

| **Recruitment** |  |  |  |  |  |  |
| --- | --- | --- | --- | --- | --- | --- |
|  | **Median** | **Mean** | **Std Dev** | **Kurtosis** | **Skewness** | **Range** |
| **Untreated** | 0.2846 | 0.2829 | 0.1424 | 3.9396 | 0.9984 | 0.6456 |
| **SMIFH2** | 0.0816 | 0.0949 | 0.0390 | 2.1311 | 0.6470 | 0.1180 |
| **CK666** | 0.0724 | 0.0846 | 0.0405 | 3.1676 | 0.9705 | 0.1569 |
| **CK+SMI** | 0.0149 | 0.0170 | 0.0109 | 4.5173 | 1.4111 | 0.0437 |

| **Turnover** |  |  |  |  |  |  |
| --- | --- | --- | --- | --- | --- | --- |
|  | **Median** | **Mean** | **Std Dev** | **Kurtosis** | **Skewness** | **Range** |
| **Untreated Fast** | 0.0957 | 0.1031 | 0.0487 | 2.6266 | 0.4639 | 0.1993 |
| **Untreated Slow** | 0.0095 | 0.0134 | 0.0146 | 4.0827 | 1.3528 | 0.0606 |
| **SMIFH2 Fast** | 0.0257 | 0.0310 | 0.0175 | 3.8492 | 1.0379 | 0.0682 |
| **SMIFH2 Slow** | 0.0087 | 0.0111 | 0.0102 | 6.8027 | 2.0662 | 0.0392 |
| **CK666 Fast** | 0.0307 | 0.0290 | 0.0167 | 2.4373 | 0.4987 | 0.0627 |
| **CK666 Slow** | 0.0061 | 0.0057 | 0.0044 | 1.9520 | -0.0906 | 0.0154 |
| **CK+SMI Fast** | 0.0034 | 0.0047 | 0.0040 | 4.0058 | 1.3056 | 0.0152 |
| **CK+SMI Slow** | 0.0046 | 0.0081 | 0.0086 | 4.1597 | 1.4876 | 0.0292 |

Interestingly, the fast turnover phase of actin kinetics shares many features in common with the recruitment phase (Fig. 1E, left). Inhibition of formin or Arp2/3 reduced the median turnover rate by 3-4 fold compared to mock treatment (Mock=0.096 fold/sec, SMIFH2=0.026 fold/sec, CK666=0.031 fold/sec) (Fig. 1E, Table 1), which is comparable to the reduction observed in the recruitment phase of actin. Moreover, inhibition of formin or Arp2/3 also altered the turnover profile of actin, generating a narrower range of turnover rates (Table 1). Given that recruitment and turnover are comparably altered by inhibition of formin or Arp2/3, our data imply a link between these phases such that rapid/slowed recruitment is paired with rapid/slowed turnover. This is also true for when formin and Arp2/3 are simultaneously inhibited, which reduces the turnover rate of actin by over 10-fold (CK+SMI=0.005 fold/sec) compared to mock treatment. In contrast, slow turnover rates exhibited only minor variations between conditions, even in circumstances where both formin and Arp2/3 are inhibited (Fig. 1E, right). To confirm that comparisons derived from the violin plots are reflective of the overall kinetics for each recruitment condition, we performed a linear regression analysis on the mean rate of recruitment and turnover for each phase (Fig. S3). We noted that the trends observed in violin plots (Fig. 1D,E) were also observed in the averaged rate slopes for all conditions tested (Fig. S3A), indicating that differences between individual rate distributions are reflective of the overall kinetics for each condition. In sum, our data indicate that inhibition of host actin nucleators substantially impairs rapid actin recruitment and turnover at the site of invasion, while only having a minor effect on the slow, lingering phase of actin turnover.

Finally, we wanted to evaluate whether the trends observed in actin recruitment and turnover at the population level were preserved at the single EB level. First, we considered whether the individual rates of actin recruitment and turnover were influenced by the net quantity of actin recruited. To do this, we plotted the rate of recruitment and turnover for each event against its maximal fold recruitment (Fig. S4A) and observed that there was no strong correlation between these traits. As such, we report that variations in recruitment and turnover rates, either within or between conditions, are not simply due to variations in net actin recruitment. Next, we wanted to determine whether the link between rapid/slowed recruitment and rapid/slowed turnover observed at the population level was also present at the level of individual EBs. To do this, we plotted the rate of recruitment for each event against its fast turnover rate (Fig. S4A) and surprisingly found that there was no correlation between the recruitment rate and turnover rate of individual EBs, indicating a level of independence between the kinetics of recruitment and turnover. One possible explanation is that predetermining factors variably influence the recruitment and turnover rates of actin at each invasion event, such that actin dynamics exhibit variability at the individual level but are not apparent when analyzed at the population level. These factors could be the local concentration of invasion-relevant translocated effectors, the density of pre-formed actin networks which serve as sites for additional actin recruitment, or the abundance of signaling co-receptors, etc. Notwithstanding, we conclude that optimal kinetics of actin recruitment are achieved when both formin and Arp2/3 are active, suggesting that *Chlamydia* relies on the nucleation of both branched and filamentous actin for optimal actin remodeling. Additionally, our data highlight the importance of quantitatively analyzing individual invasion events.

### Formin and Arp2/3 are collaboratively recruited to the site of invasion

The observation that formin and Arp2/3 promote actin recruitment during invasion raise the possibility that these nucleators act collaboratively, thereby enhancing the capacity of *Chlamydia* to generate actin-rich invasion structures. If nucleator collaboration does indeed play a role in invasion, we expect that both formin and Arp2/3 will be present within a single invasion event, and that inhibiting either nucleator will impair recruitment of its collaboration partner. Although multiple species of formin were recruited during invasion (Fig. 1A), we chose to monitor Fmn1 since this protein was specifically identified in a prior siRNA screen of cytoskeletal effectors with a role in *Chlamydia* invasion (35). We observed that GFP-Fmn1 was robustly recruited during invasion and experienced a substantial defect in recruitment upon pharmacological inhibition of Arp2/3 (Fig. 2A). Likewise, GFP-Arp3, which has been validated as a marker for the Arp2/3 complex (38), was robustly recruited during invasion and also demonstrated reduced recruitment following inhibition of formin (Fig. 2B). Altogether, this indicates that recruitment of Fmn1 and Arp2/3 are reciprocally enhanced by their respective actin nucleating functions and subsequent actin polymerization.

**Figure 2:**
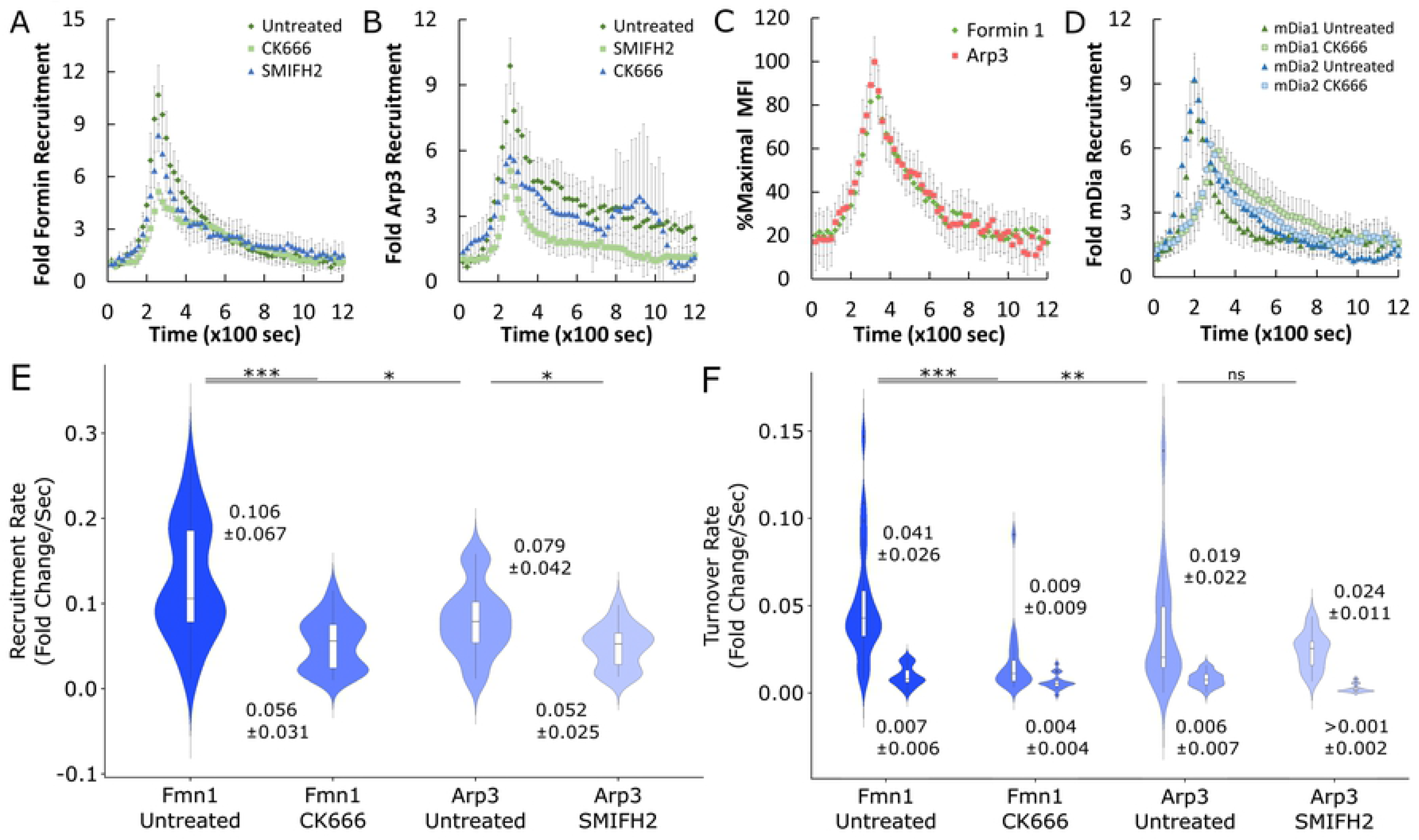
Formin 1 and Arp3 collaboration enhances recruitment kinetics at entry sites. (A,B) Cos7 cells were transfected with (A) GFP-formin 1 isoform 1B (GFP-Fmn1) or (B) GFP-Arp3 for 24 hours prior to mock-treatment or pretreatment with 10µM SMIFH2 or 100µM CK666 for 1 hour. Quantitative live-cell imaging of invading CMTPX- labeled *Chlamydia* (MOI=20) was performed as described in Fig. 2B. Briefly, images were acquired once every 20 seconds for 30 minutes and assembled into videos. MFI of recruitment events were obtained, and background fluorescence subtracted, before quantifying and plotting the fold recruitment values +/- SEM of Fmn1 or Arp3 for each timepoint. Data was compiled from a minimum N=16 recruitment events. (C) Cos7 cells were co-transfected with GFP- Fmn1 and mCherry-Arp3 for 24 hours prior to infection with unlabeled *C. trachomatis* (MOI=20) followed by quantitative live-cell imaging of invasion. Image acquisition and processing was performed as described above with a few modifications. Since CMTPX and mCherry-Arp3 fluorescence overlap, unlabeled *Chlamydia* was used, thus recruitment events are defined simply as regions containing elevated GFP-Fmn1 fluorescence compared to local background. MFI of both GFP-Fmn1 and mCherry-Arp3 were obtained from the recruitment ROI, and background was subtracted from both GFP and RFP channels independently. Maximal MFI was derived from both channels and used to normalize fluorescence as percent maximal MFI for each protein. Normalized MFI values for each timepoint were compiled from 33 events, plotting mean %Max MFI +/- SEM. (D) Recruitment of mEmerald-mDia1 and mEmerald-mDia2 was monitored in the presence and absence of 100µM CK666 using the method described above for monitoring the recruitment of GFP-Fmn1 and GFP-Arp3. Data is displayed as mean fold recruitment for each timepoint +/- SEM compiled from a minimum N=22 recruitment events. (E,F) Kinetics of Fmn1 and Arp3 recruitment and turnover were analyzed in the presence and absence of 100µM CK666 or 10µM SMIFH2, respectively, using the same methodology described in Fig. 1D, E. Violin plots contain a minimum N=16 individual rates, reporting the median rate +/- SD. Statistical significance for violin plots was determined by Kolmogorov-Smirnov test. All data are representative of at least 3 independent experiments, * P ≤ 0.05, ** P ≤ 0.01, *** P ≤ 0.001.

Next, we determined whether Fmn1 and Arp2/3 are co-recruited within individual invasion events, thus providing the necessary context for nucleator collaboration. To do this, we co-expressed GFP- Fmn1 and mCherry-Arp3 and measured the recruitment of both proteins during invasion by time-lapse imaging followed by fluorescence quantification (Fig. 2C). We observed that all events we monitored exhibited both Fmn1 and Arp3 accumulation, indicating that they are co-recruited by *Chlamydia*. To measure the timing of their arrival, we normalized the intensity of recruitment to the maximal MFI of each protein at each timepoint, expecting that any delays in protein recruitment would be represented by a shift in their recruitment profile. In contrast, we noted that the recruitment profiles of both Fmn1 and Arp3 were remarkably similar and achieved maximal MFI at the same timepoint, indicating that both proteins arrived at the entry site within 20 seconds of each other, the temporal resolution of our imaging protocol. Given both the simultaneous recruitment of Fmn1 and Arp2/3 and the reciprocal dependence of their respective activities for optimal recruitment, we conclude that Fmn1 and Arp2/3 collaborate during invasion to promote actin recruitment. This observation is particularly novel, given that there are no other instances of the utilization of Fmn1 by bacterial pathogens. Lastly, given the robust recruitment of Fmn1, we were interested in whether other formin species such as DRFs, which require upstream activation by Rho-GTPases, could also participate in nucleator collaboration. To this end, we found that mEmerald-mDia1 and mEmerald-mDia2 were also recruited during invasion, and that their recruitment was impaired by Arp2/3 inhibition (Fig. 2D), suggesting that both DRF and non-DRF formins collaborate with Arp2/3 during invasion.

### Recruitment and turnover profiles of Fmn1 and Arp2/3 are altered by reciprocal inhibition

Using the same methodology to evaluate actin kinetics (Fig. 1D,E), we compared the recruitment and turnover profiles of formin and Arp2/3 to more closely evaluate how inhibition of each protein alters their recruitment and turnover dynamics during invasion. We noted that in absence of inhibitor, Fmn1 recruitment was slightly faster than Arp3, exhibiting a median recruitment rate that was 25 percent higher than Arp3 (Fmn1=0.106 fold/sec vs. Arp3=0.079 fold/sec) (Fig. 2E, Table 2). However, this difference was not preserved in the averaged rate slopes of Fmn1 and Arp3 recruitment (Fig. S3B), indicating that while their rate distributions differ, their overall kinetics are comparable. Upon reciprocal inhibition, the median recruitment rates for both Fmn1 and Arp3 decreased by 47 percent and 34 percent, respectively, resulting in a comparable recruitment rate for both nucleators (Fmn1=0.056 fold/sec vs. Arp3=0.052 fold/sec) (Fig. 2E, Table 2). Together, this indicates that collaboration between the activities of Fmn1 and Arp3 not only enhance the rate of actin recruitment (Fig. 1D), but also reciprocally enhance their respective recruitment rates. Although the recruitment profiles of Fmn1 and Arp3 were largely comparable, we found that their fast turnover profiles exhibited substantial differences. In particular, we observed that mock-treated Fmn1 achieved a median fast turnover rate that was over twofold higher than that of mock-treated Arp3 (Fmn1=0.041 fold/sec vs. Arp3=0.019 fold/sec) (Fig. 2F, Table 2). Furthermore, while CK666 treatment reduced the turnover rate of Fmn1 by 4-fold compared to mock-treatment (Mock=0.041 fold/sec vs. CK666=0.009 fold/sec), SMIFH2 treatment did not significantly alter the rate of Arp3 turnover (Mock=0.019 fold/sec vs. SMIFH2=0.024 fold/sec).

**Table 2.** Statistical values associated with Fmn1 and Arp2/3 recruitment and turnover in presence and absence of inhibitor.

| Recruitment |  |  |  |  |  |  |
| --- | --- | --- | --- | --- | --- | --- |
|  | Median | Mean | Std Dev | Kurtosis | Skewness | Range |
| Fmn1 Untreated | 0.1056 | 0.1252 | 0.0666 | 2.1733 | 0.3321 | 0.2530 |
| Fmn1 CK666 | 0.0556 | 0.0529 | 0.0306 | 1.9177 | 0.2206 | 0.1054 |
| Arp3 Untreated | 0.0786 | 0.0801 | 0.0416 | 2.4882 | 0.2980 | 0.1466 |
| Arp3 SMIFH2 | 0.0524 | 0.0492 | 0.0252 | 2.1480 | 0.0563 | 0.0844 |

| Turnover |  |  |  |  |  |  |
| --- | --- | --- | --- | --- | --- | --- |
|  | Median | Mean | Std Dev | Kurtosis | Skewness | Range |
| Fmn1 Untreated Fast | 0.0409 | 0.0444 | 0.0255 | 3.2119 | 0.8429 | 0.0995 |
| Fmn1 Untreated Slow | 0.0069 | 0.0083 | 0.0058 | 2.2052 | 0.4465 | 0.0206 |
| Fmn1 CK666 Fast | 0.0093 | 0.0120 | 0.0089 | 3.1418 | 1.0889 | 0.0313 |
| Fmn1 CK666 Slow | 0.0043 | 0.0051 | 0.0041 | 3.5222 | 0.8693 | 0.0175 |
| Arp3 Untreated Fast | 0.0187 | 0.0278 | 0.0216 | 2.4568 | 0.7775 | 0.0771 |
| Arp3 Untreated Slow | 0.0064 | 0.0068 | 0.0042 | 2.2873 | 0.2086 | 0.0152 |
| Arp3 SMIFH2 Fast | 0.0240 | 0.0228 | 0.0106 | 2.2091 | 0.0892 | 0.0368 |
| Arp3 SMIFH2 Slow | 0.0004 | 0.0015 | 0.0023 | 3.3368 | 1.2892 | 0.0076 |

The dissimilarity between the recruitment and turnover phases of Fmn1 and Arp3 suggest that they are governed by different mechanisms, an assertion that is further supported by our observation that there is no correlation between the rate of recruitment and turnover for individual EBs regarding Fmn1 (R^2^=0.1345) or Arp3 (R^2^=0.1561). Additionally, while we observed a moderate correlation between the maximal fold recruitment and the recruitment rate of Arp3 (R^2^=0.621), all other conditions had no correlation (Fig. S4B). In sum, these data suggest that Fmn1 and Arp2/3 collaboration is necessary for rapid turnover of Fmn1, but not Arp2/3. While we observed some slight differences in the slow turnover phases of Fmn1 and Arp3 (Fig. 2F), we noted that the variability of these data are quite high. As such, it is difficult to give a precise account of how reciprocal inhibition affects the residual turnover of Fmn1 and Arp3, although the miniscule turnover rates within this phase suggest that its contribution to the overall turnover of Fmn1 and Arp2/3 is minor. Finally, we noted that the recruitment and turnover kinetics of mDia1 and mDia2 were comparable to Fmn1 both in mock- and CK666-treated groups (Fig. S5, Table S1), indicating that all formin species are regulated comparably during invasion. Altogether, we conclude that kinetics of recruitment and turnover of Fmn1 and DRFs are enhanced by collaboration with Arp2/3, and reflect the kinetics observed for actin recruitment and turnover (Fig. 1D,E). In contrast, nucleator collaboration enhances recruitment of Arp3, but does not affect its turnover, suggesting that the factors which dictate turnover of Fmn1 and Arp3 differ in some respect. The difference could be accounted for by the stability of the actin network serving as scaffold for either Fmn1 or Arp2/3. Thus, the actin network produced by Fmn1 and Arp3 collaboration likely serves as a binding site which enhances Fmn1 and Arp2/3 recruitment, while having varied effects on their turnover.

### Deletion of TarP results in defective invasion and dramatically alters kinetics of actin recruitment and turnover

Thus far we have demonstrated that *C. trachomatis* invasion relies on the recruitment and actin nucleating activities of the host formins Fmn1, mDia1, and mDia2, while specifically identifying a collaboration between Fmn1 and Arp2/3 to enhance actin remodeling. It was previously demonstrated that the *Chlamydia* effector TarP not only binds filamentous actin directly *in vitro*, but also recruits Arp2/3 via signaling through the Arp2/3 activator WAVE2 (24,39). Furthermore, it was recently established that deletion of TarP resulted in a significant defect in pathogen internalization (40). Given the central role of TarP in coordinating internalization of *Chlamydia* by manipulation of the host actin cytoskeleton (13), we sought to determine whether TarP deletion prevented actin recruitment. Since the TarP deletion strain (ΔTarP) expresses GFP as a byproduct of the homologous recombination event used to create the TarP deletion via FRAEM, we utilized an RFP-conjugated form of actin (mRuby-LifeAct). We observed that actin recruitment was attenuated and more diffuse in ΔTarP than actin recruited by a strain in which TarP was restored by *cis*-complementation (*cis*-TarP) (Fig. 3A). To determine if attenuated actin recruitment is due to restricted utilization of Fmn1 and Arp2/3, we tested whether deletion of TarP rendered *Chlamydia* unresponsive to the subsequent application of inhibitors to formin or Arp2/3 (Fig. 3B). In agreement with previous findings, ΔTarP exhibited a substantial impairment to invasion efficiency relative to *cis*-TarP (40). Contrary to our expectations, inhibition of formin or Arp2/3 resulted in a further reduction in pathogen internalization, however, invasion efficiency was not substantially worsened by simultaneous inhibition. This suggests that while the invasion-associated activities of formin and Arp2/3 are not entirely TarP dependent, deletion of TarP renders *Chlamydia* invasion more sensitive to formin and/or Arp2/3 inhibitors.

**Figure 3:**
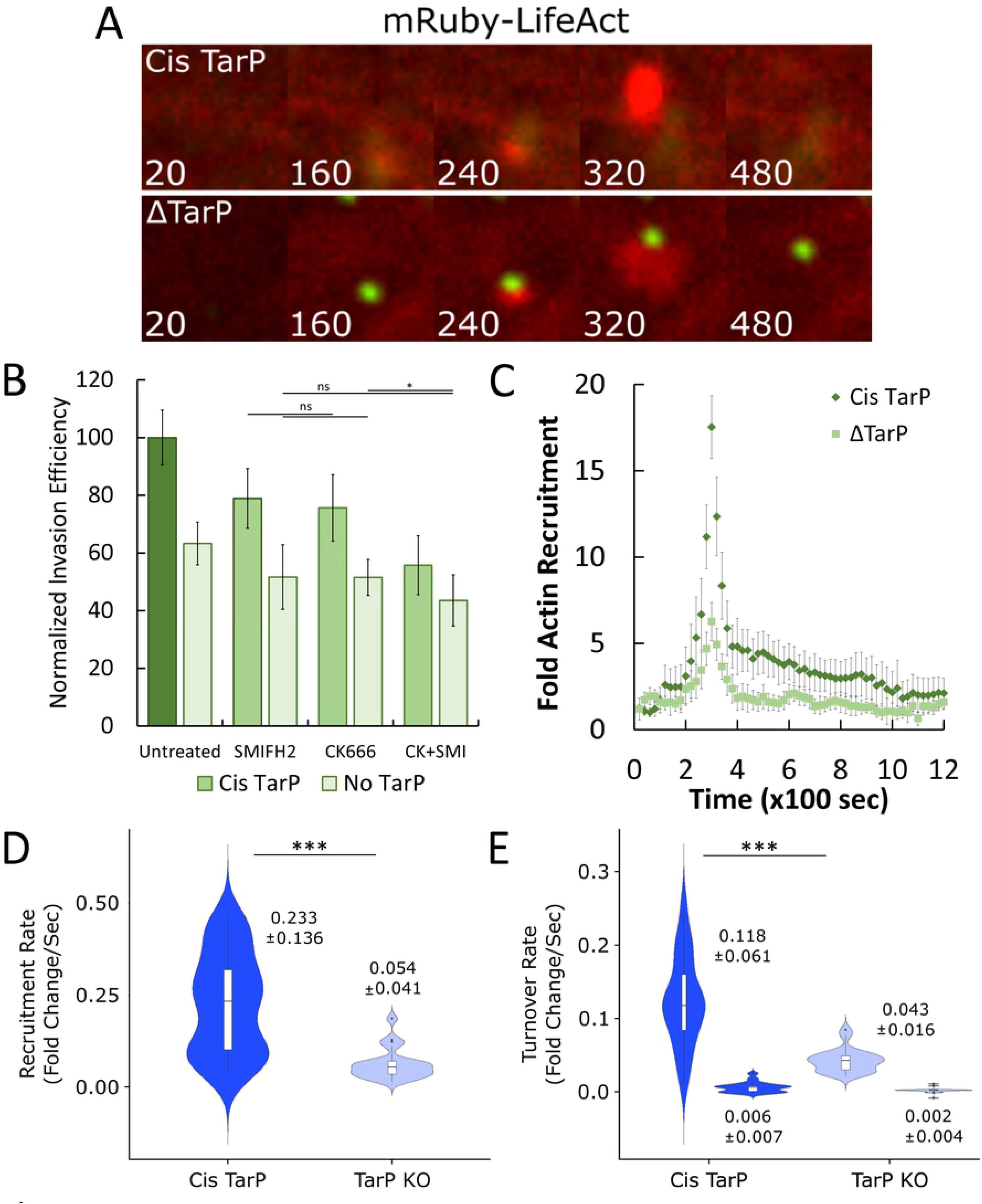
Deletion of TarP impairs invasion and attenuates kinetics of actin recruitment and turnover. (A) Cos7 cells were transfected with mRuby-LifeAct for 24 hours prior to infection with either a TarP deletion mutant of *C. trachomatis* (ΔTarP) or a strain in which TarP was restored by *cis*-complementation (*cis*-TarP) at MOI=20. Infection was monitored by live-cell confocal microscopy, obtaining images every 20 seconds for 30 minutes to identify sites exhibiting actin recruitment. (B) Cos7 cells were mock-treated or pretreated with inhibitors against formin (10µM SMIFH2), Arp2/3 (100µM CK666) or both (CK+SMI) for 1hr. Cells were infected with either ΔTarP or *cis*-TarP at MOI=50 and stained using the “in-and-out” method described in Fig. 1B. Results were normalized against mean invasion efficiency of *cis*-TarP in mock treated cells (*cis*-TarP untreated mean = 52% invasion efficiency) and plotted as normalized mean +/- SEM. Data was collected from 12 fields, with each field containing an average of 127 Chlamydiae. Statistical significance was determined by T- test. (C) Cos7 cells were transfected with mRuby-LifeAct for 24 hours prior to infection with either *cis*-TarP or ΔTarP at MOI=20. Quantitative live-cell imaging of invading *Chlamydia* was performed as described in Fig. 1C. Briefly, images were acquired once every 20 seconds for 30 minutes and assembled into videos. MFI of recruitment events were obtained, and background fluorescence subtracted, before quantifying and plotting the fold recruitment values +/- SEM of mRuby- LifeAct for each timepoint compiled from a minimum N=22 recruitment events. (D,E) Kinetics of mRuby-LifeAct recruitment and turnover were analyzed for *cis*-TarP and ΔTarP using the same methodology described in Fig. 1D,E. Violin plots contain a minimum N=22 individual rates, reporting the median rate +/- SD. Statistical significance was determined by Kolmogorov-Smirnov test. All data are representative of at least 3 independent experiments, * P ≤ 0.05, *** P ≤ 0.001.

To more precisely evaluate the effect of TarP deletion on actin recruitment, we tracked the accumulation of mRuby-LifeAct at *Chlamydia* entry sites and analyzed the kinetics of actin recruitment and turnover (Fig. 3 C-E). We observed that *cis*-TarP robustly recruited actin, resulting in a 3-fold increase in actin accumulation compared to ΔTarP (Fig. 3C). We noted that *cis*-TarP featured a median actin recruitment rate that is comparable to wild-type *C. trachomatis* (*cis*-TarP=0.233 fold/sec vs. wild-type=0.285 fold/sec) (Fig. 1D), while loss of TarP reduced the rate of actin recruitment by 4-fold (ΔTarP=0.054 fold/sec) (Fig. 3D, Table 3). Likewise, TarP deletion slowed the rate of actin turnover by nearly 3-fold (*cis*-TarP=0.117 fold/sec vs. ΔTarP=0.043 fold/sec) (Fig. 3E), indicating that TarP contributes not only to the rapid recruitment of actin, but also assists in facilitating rapid turnover. We did not observe any correlation between the rate of recruitment and turnover in *cis*-TarP (R^2^=0.115) and weak correlation in ΔTarP (R^2^=0.287), suggesting that TarP alone is insufficient to explain the independent regulation of actin recruitment and turnover (Fig. S6A). However, TarP deletion also alters the slow turnover phase of actin (Fig. 3E), eliminating the boundary between rapid and lingering actin turnover that is readily observable in both *cis*-TarP and wild-type strains. Taken together, we conclude that while TarP is insufficient as a sole means of coordinating actin turnover, it is clear that the effector is involved in the process in some capacity. Thus, we propose that TarP may be a necessary component for rapid actin turnover, serving as a signaling nexus for both actin polymerizing and actin turnover machinery.

**Table 3.**
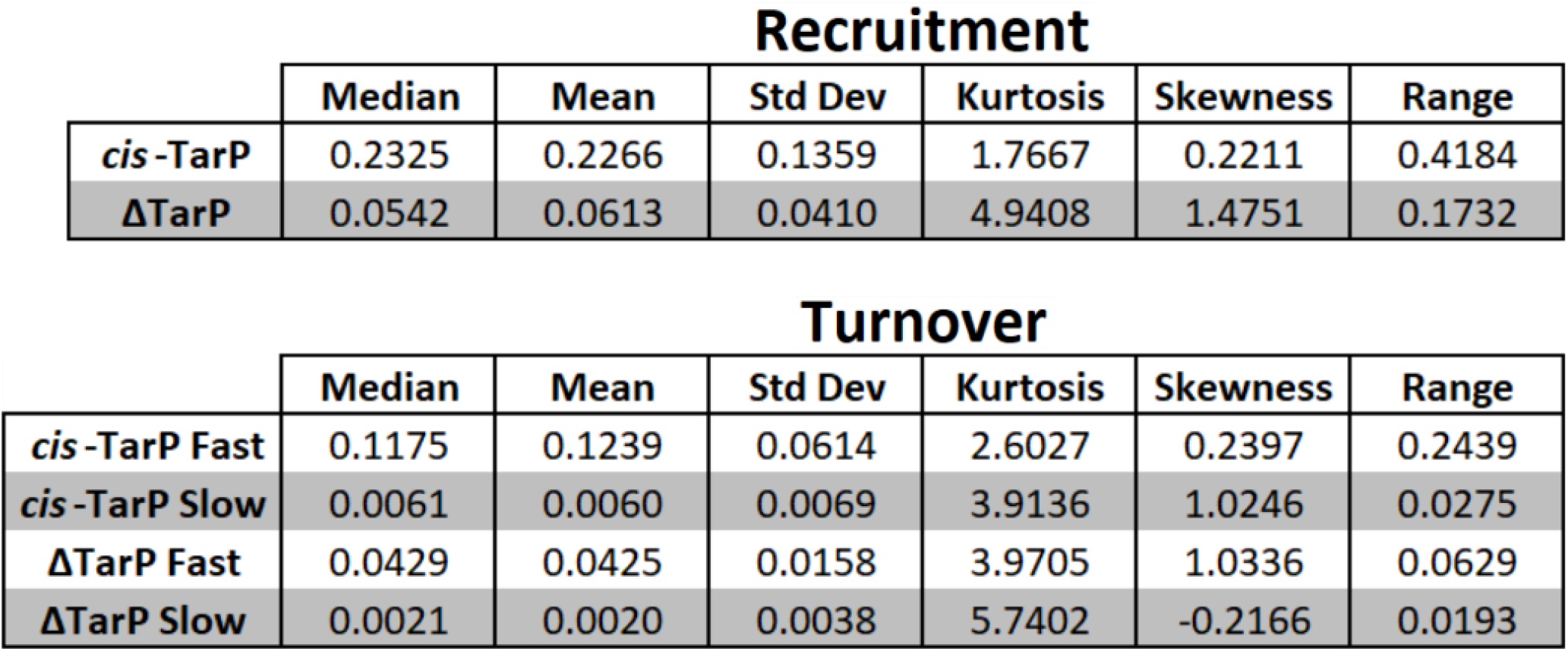
Statistical values associated with actin recruitment and turnover in ΔTarP and *cis*-TarP backgrounds.

### Deletion of TarP prevents recruitment of Fmn1 and attenuates kinetics of Arp3 recruitment and turnover

Next, we determined whether the impaired recruitment kinetics of actin in ΔTarP *Chlamydia* are due to deficient recruitment of Fmn1 and Arp3 by tracking mCherry-conjugated forms of both proteins at entry sites (Fig. 4A-C). While *cis*-TarP robustly recruited both Fmn1 and Arp3, ΔTarP experienced attenuated Arp3 recruitment and failed to recruit Fmn1 altogether (Fig. 4A). Quantitative imaging of Fmn1 and Arp3 recruitment revealed that Arp3 recruitment in ΔTarP was 2-fold less robust than *cis*-TarP, while only recording trace signals of Fmn1 recruitment in a ΔTarP background (Fig. 4B,C). However, since ΔTarP bacteria are highly fluorescent, it is likely that the trace RFP signal is due to fluorescence bleed-through rather than bona-fide recruitment. Collectively, these data indicate that while Fmn1 recruitment is TarP-dependent, the recruitment of Arp2/3 could be driven in part by other factors present during invasion such as host receptor activation or other *Chlamydia* effectors (e.g., TmeA) (41,42).

**Figure 4:**
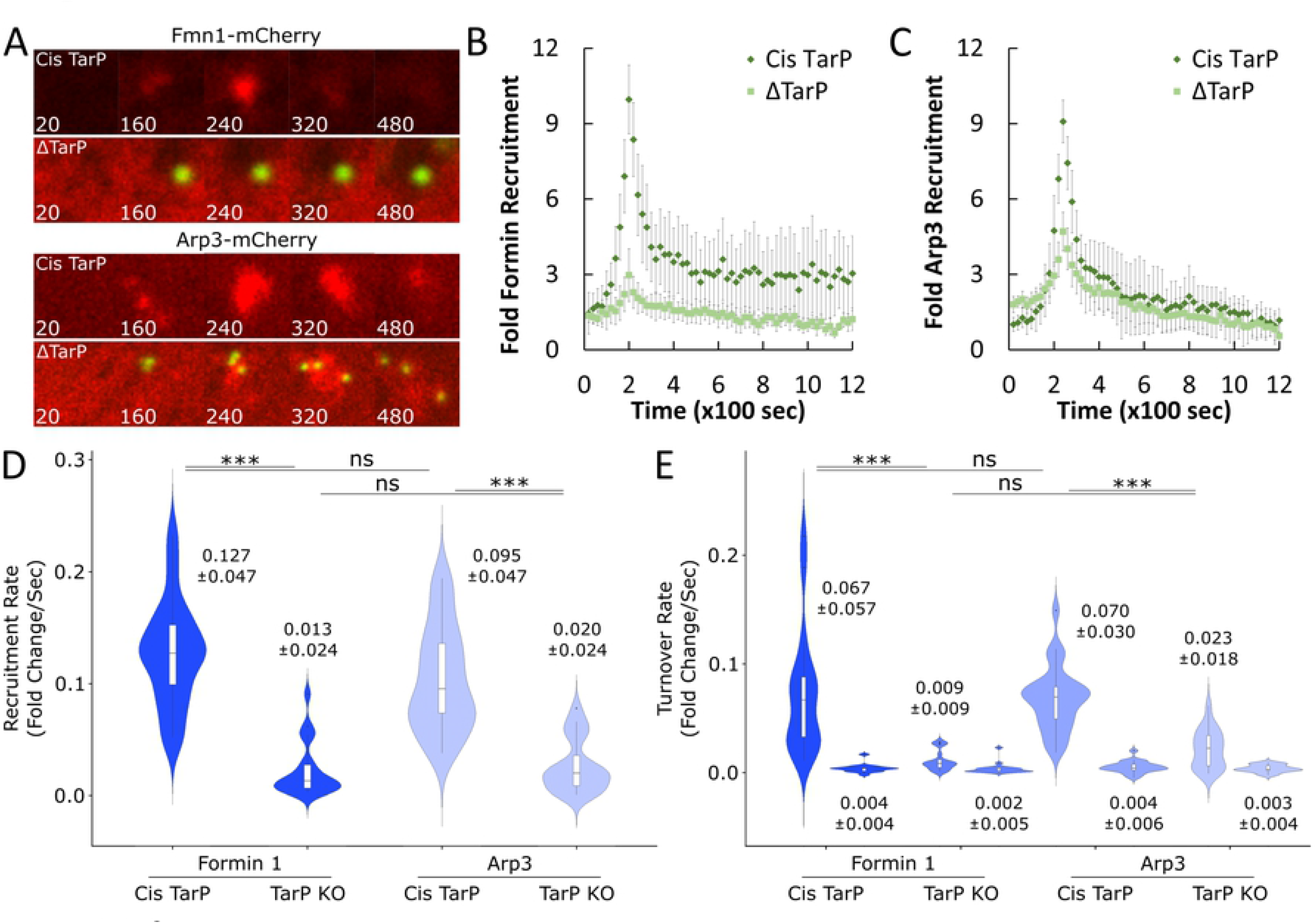
Deletion of TarP attenuates kinetics of Fmn1 and Arp3 recruitment and turnover. (A) Cos7 cells were transfected with Fmn1-mCherry or Arp3-mCherry for 24 hours prior to infection with either ΔTarP or *cis*-TarP at MOI=20. Infection was monitored by live-cell confocal microscopy, obtaining images every 20 seconds for 30 minutes to identify sites exhibiting Fmn1 or Arp3 recruitment. (B,C) Cos7 cells were transfected with (B) Fmn1-mCherry or (C) Arp3-mCherry for 24 hours prior to infection with either *cis*-TarP or ΔTarP at MOI=20. Quantitative live-cell imaging of invading *Chlamydia* was performed as described in Fig. 1C. Briefly, images were acquired once every 20 seconds for 30 minutes and assembled into videos. MFI of recruitment events were obtained, and background fluorescence subtracted, before quantifying and plotting the fold recruitment values +/- SEM of Fmn1 or Arp3 for each timepoint compiled from a minimum N=18 recruitment events. (D,E) Kinetics of Fmn1-mCherry or Arp3-mCherry recruitment and turnover were analyzed for *cis*-TarP and ΔTarP using the same methodology described in Fig. 1D,E. Violin plots contain a minimum N=18 individual, reporting the median rate +/- SD. Statistical significance was determined by Kolmogorov-Smirnov test. All data are representative of at least 3 independent experiments, *** P ≤ 0.001.

To better understand the role of TarP in the recruitment of host actin nucleators, we analyzed the kinetics of Fmn1 and Arp3 recruitment and turnover following TarP deletion. We noted that loss of TarP slowed the recruitment of Fmn1 by 10-fold (*cis*-TarP=0.127 fold/sec vs. ΔTarP=0.013 fold/sec) and Arp3 by 5-fold (*cis*-TarP=0.095 fold/sec vs. ΔTarP=0.020 fold/sec) (Fig. 4E, Table 4). By comparison, reciprocal inhibition of Fmn1 or Arp3 slowed the recruitment of either nucleator by just over 2-fold (Fig. 2E). Additionally, we noted that the rate of Fmn1 recruitment in a ΔTarP background was nearly negligible; barring outliers, the median recruitment rate clustered around 0.0093 fold/sec, which is consistent with our earlier assessment that Fmn1 is likely not recruited in the absence of TarP (Fig. 4A,B). In sum, these data indicate that TarP contributes to the recruitment of Fmn1 and Arp3, and that loss of TarP significantly attenuates the recruitment of both nucleators.

**Table 4.** Statistical values associated with Fmn1 and Arp2/3 recruitment and turnover in ΔTarP and *cis*-TarP backgrounds.

Recruitment
|  | Median | Mean | Std Dev | Kurtosis | Skewness | Range |
| --- | --- | --- | --- | --- | --- | --- |
| Fmn1 *cis*-TarP | 0.1270 | 0.1274 | 0.0472 | 2.9163 | 0.3413 | 0.1800 |
| Fmn1 ΔTarP | 0.0129 | 0.0222 | 0.0240 | 4.4560 | 1.5554 | 0.0879 |
| Arp3 *cis*-TarP | 0.0952 | 0.1032 | 0.0465 | 2.1808 | 0.4840 | 0.1562 |
| Arp3 ΔTarP | 0.0201 | 0.0262 | 0.0238 | 2.5129 | 0.8832 | 0.0773 |

Turnover
|  | Median | Mean | Std Dev | Kurtosis | Skewness | Range |
| --- | --- | --- | --- | --- | --- | --- |
| Fmn1 *cis*-TarP Fast | 0.0670 | 0.0721 | 0.0568 | 4.0345 | 1.3100 | 0.2062 |
| Fmn1 *cis*-TarP Slow | 0.0037 | 0.0040 | 0.0041 | 7.3170 | 1.8364 | 0.0188 |
| Fmn1 ΔTarP Fast | 0.0086 | 0.0110 | 0.0091 | 2.5094 | 0.9283 | 0.0277 |
| Fmn1 ΔTarP Slow | 0.0022 | 0.0037 | 0.0051 | 12.1221 | 3.0129 | 0.0236 |
| Arp3 *cis*-TarP Fast | 0.0696 | 0.0682 | 0.0296 | 3.6494 | 0.6393 | 0.1309 |
| Arp3 *cis*-TarP Slow | 0.0041 | 0.0051 | 0.0054 | 4.1060 | 0.6582 | 0.0252 |
| Arp3 ΔTarP Fast | 0.0225 | 0.0217 | 0.0178 | 2.3832 | 0.4634 | 0.0627 |
| Arp3 ΔTarP Slow | 0.0029 | 0.0038 | 0.0035 | 2.1958 | 0.0188 | 0.0121 |

Additionally, the fast turnover rates for Fmn1 and Arp3 in *cis*-TarP (Fmn1=0.067 fold/sec, Arp3=0.070 fold/sec) were 3-6 fold faster than those observed in ΔTarP (Fmn1=0.009 fold/sec, Arp3=0.023 fold/sec) (Fig. 4E, Table 4), suggesting that TarP also contributes to rapid turnover of Fmn1 and Arp3. In contrast, the slow turnover rates of Fmn1 and Arp3 were mostly unaffected by TarP deletion, suggesting that TarP predominantly affects the rapid turnover phase of formin and Arp2/3 (Fig. 4E). Since the turnover of Fmn1 and Arp2/3 (Fig. 2F) is linked to actin turnover (Fig. 1E), the role of TarP as a determinant of Fmn1 and Arp2/3 turnover kinetics is likely related to its effects on the turnover of the actin network assembled post-recruitment (Fig. 3), and to which these nucleators bind. This is further supported by the fact that recruitment and turnover rates are not correlated in Fmn1 (R^2^=0.111), Arp3 (R^2^<0.001), or actin (R^2^=0.115) (Fig. S6), which indicates that the recruitment and turnover phases for all three proteins are independently regulated. Altogether, we conclude that TarP is necessary for the establishment of a robust actin network comprised of branched and filamentous actin, which contributes to the rapid recruitment and turnover of Fmn1 and Arp3.

Taken together, our data establish TarP as a hub for actin polymerization, particularly in its ability to drive the recruitment of nucleators for both branched and filamentous actin. As such, *Chlamydia* utilizes TarP to create the necessary conditions which promote Fmn1 and Arp2/3 collaboration, which the pathogen exploits to generate a robust actin network at the site of entry. Inhibition of either Fmn1 or Arp2/3 has severe negative effects on *Chlamydia* invasion, and results in defective recruitment of actin, Fmn1 and Arp3. Finally, our data implicate turnover of actin and its effectors as a necessary component of efficient invasion, as TarP deletion or inhibitor treatment both impair pathogen internalization and restrict turnover of actin, Fmn1 and Arp3. Thus, we establish that TarP is not only a platform for the recruitment of nucleators for both branched and filamentous actin networks, but likely also contributes to actin turnover during *Chlamydia* internalization.

## Discussion

In this study, we identified a signature actin dynamics associated with efficient invasion that was largely determined by the combined signaling functions of TarP, Arp2/3, and Fmn1. We demonstrated a collaborative role for two actin nucleators, formin and Arp2/3, which *Chlamydia* exploits to generate a robust actin modulatory network at pathogen entry sites to promote its internalization. By monitoring the kinetics of recruitment, we highlighted the role of formin and Arp2/3 collaboration in accelerating the rate of actin recruitment, as well as reciprocally enhancing the recruitment of actin nucleators. Furthermore, we emphasized the necessity for actin turnover in order to achieve pathogen internalization and introduced a role for TarP as a coordinator of nucleator recruitment and actin turnover.

### Actin dynamics at the site of chlamydial invasion

Intracellular pathogens often target the actin cytoskeleton during entry, leading to robust actin recruitment and subsequent formation of invasion-associated structures. Relative to *Chlamydia*, actin recruitment in other intracellular bacteria is longer lived (17,43), lasting on the order of 5-15 minutes; in contrast, we have shown that *Chlamydia* actin recruitment is both robust and transient, emphasizing the importance of rapid actin accumulation and turnover during its invasion. To our knowledge, no equivalent study on the rate of actin recruitment during the internalization of an intracellular pathogen has been conducted, thus a direct comparison on the kinetics of actin recruitment between pathogens is not possible. As such, we believe our study provides a valuable new approach that could be used in future studies to improve contextualization of protein dynamics for other intracellular pathogens in addition to *Chlamydia*.

Using these tools, we found that actin is recruited by *Chlamydia* at a rapid pace, owing in part to the collaboration between Fmn1 and Arp2/3. One explanation for the high rate of recruitment observed in our study is the establishment of a positive feedback loop, where actin branching by Arp2/3 and filament elongation by Fmn1 create additional nodes for Arp2/3 and Fmn1 to bind (Fig. 5C). Iterations of this process would account for both the high recruitment rate of actin in addition to providing an explanation for why inhibition of Fmn1 and Arp2/3 comparably affect the recruitment rate of actin. However, this model does not immediately make apparent why actin recruitment is transient, nor does it provide explanation for the slow and persistent actin recruitment observed following inhibition of both formin and Arp2/3. Other host actin nucleators such as SPIRE, APC, or Cordon-bleu have not yet been identified in siRNA screens of invasion-associated proteins, nor has any evidence of their involvement in *Chlamydia* invasion been given (44). Although TarP has been shown via pyrene assay to serve as an actin nucleator, it remains unknown whether it also performs this function in cell lines or *in-vivo* (45). However, based on results described here, TarP deletion drastically impaired the kinetics of recruitment and turnover of both actin and its nucleators, indicating that actin nucleation is predominantly driven by host nucleators rather than TarP itself.

**Figure 5:**
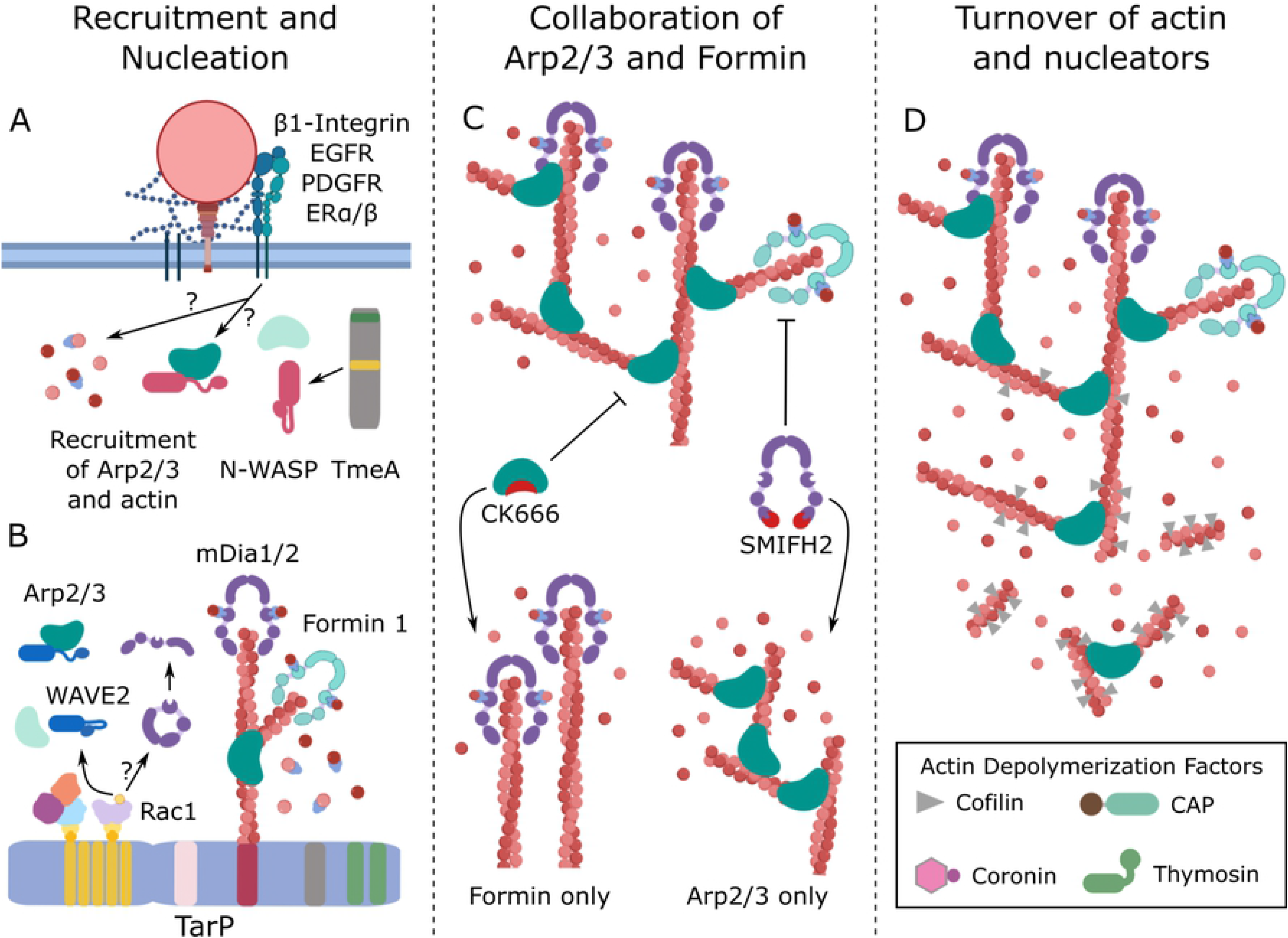
Proposed model for collaboration between formin and Arp2/3 and subsequent rapid actin recruitment and turnover. (A) *C. trachomatis* engages and activates a multitude of host receptors whose activation is linked to the recruitment of Arp2/3 and actin in non-invasion contexts. It is currently unknown whether receptor activation contributes to actin remodeling during invasion. Concurrent with receptor engagement, *Chlamydia* secretes the effectors TmeA and TarP via Type III secretion system; each effector activates Arp2/3 through N-WASP or WAVE2, respectively. (B) In addition to recruiting Arp2/3, TarP signaling through Rac1 may also function to recruit mDia1 and/or mDia2. TarP also serves as a platform for actin nucleation by direct interaction with filamentous actin, and thus may be a scaffold for the variety of recruited actin nucleators (Arp2/3, Fmn1, mDia1/2) to act upon. (C) Collaboration between Arp2/3 and formin via their respective actin branching and elongating activities increases the rate of actin recruitment. Inhibition of Arp2/3 by CK666 or formin by SMIFH2 prevents this collaboration and constrains the actin network into either filamentous-only or branching-only networks. (D) The actin network generated by *Chlamydia* is rapidly turned over alongside the actin nucleators found within it. Although the precise mechanism by which this process occurs remains unknown, four host actin depolymerization factors (ADF-Cofilin, cyclase associated protein (CAP), Coronin, and β-thymosin) have been implicated by an siRNA screen to participate in invasion.

In addition to rapid actin recruitment, we also observed a two-phase actin turnover process starting with a brief fast turnover phase (100-180 sec) followed by a prolonged slow turnover phase (400- 600 sec). The rate of turnover was attenuated by either inhibition of formin or Arp2/3, or by loss of TarP, indicating that the activity of host nucleators and TarP are associated with actin turnover. Although this process is relatively understudied in intracellular bacterial invasion, actin turnover is of clear importance for bacterial internalization, as dysregulation of actin turnover machinery commonly impairs bacterial entry. For instance, impairment or hyperactivation of the actin depolymerizing protein ADF-cofilin restricts internalization of *S. enterica*, while overexpression of the ADF-cofilin kinase LIMK reduced *L. monocytogenes* ruffle formation and internalization (46,47). In *C. burnetii* and *L. monocytogenes*, time-course experiments tracking actin recruitment also demonstrated a clear spike in actin recruitment followed by a two-phase rapid/residual turnover profile (43,47). Thus, the actin turnover profile observed in our study may simply reflect normal actin turnover dynamics, as opposed to being intrinsic to invading *Chlamydia*. Given the importance of host actin depolymerization factors like gelsolin, ADF-cofilin and their regulators for efficient internalization of other intracellular bacteria, it may be that such proteins are also responsible for actin turnover during *Chlamydia* invasion (32,48–50). Indeed, an siRNA screen of *C. muridarum* invasion-associated proteins identified four actin depolymerization factors, ADF-cofilin, cyclase associated protein (CAP), Coronin, and β-thymosin, each of which reduced bacterial entry upon silencing (Fig. 5D) (35). Thus, one avenue for future study is to characterize the role of ADF-cofilin and other actin depolymerization factors in *Chlamydia* invasion and whether dysregulation or silencing of these proteins alters the kinetics of actin turnover. Furthermore, additional study is required to determine what circumstances prompt the termination of actin recruitment and how *Chlamydia* achieves actin recruitment in absence of host nucleators. Of particular interest is the apparent correlation between actin turnover and the presence of functional TarP, Arp2/3, and Formin 1. This raises a number of possible mechanisms that trigger rapid turnover, including actin network density, physical stress that might sensitize the actin network to turnover, or the actin network having a configuration that is preferred by the local depolymerization machinery present.

### Tarp-dependent vs. TarP-independent invasion

While our data impress the importance of TarP in promoting the recruitment and turnover of Arp2/3, which occurs via signaling through the WAVE2 complex, we also observed that a portion of Arp2/3 was recruited in a TarP-independent manner. Given recent evidence demonstrating that the *Chlamydia* effector TmeA also activates Arp2/3 during invasion by signaling through N-WASP it is likely that the residual recruitment of actin and Arp2/3 in ΔTarP strains is due, at least in part, to the activity of TmeA (Fig. 5A) (41,42). A direct comparison between WT, ΔTarP, ΔTmeA, ΔTarP/ ΔTmeA strains with regards to dynamics of actin, Fmn1, and Arp2/3 at the invasion site would be required to assess accurately their relative contributions. However, some inferences can be drawn from the data reported here. We observed that the predominant fraction of Arp2/3 recruitment was linked to TarP (Fig. 4C), indicating that the contribution of TmeA in this process is auxiliary. Furthermore, invasion of TarP-deficient mutants that retained wild type TmeA was associated with a radically different actin kinetics, i.e. a slower rate of recruitment and an extremely slow turnover (Fig. 3B-E), which would be expected to be associated with slower pathogen internalization. It is unclear if Arp2/3 collaborates with Fmn1 in the context of TmeA- dependent, but TarP-independent invasion and actin dynamics, however available data indicate no collaboration. It may be that the actin network formed by the TmeA/N-WASP/Arp2/3 axis does not offer binding sites for Fmn1, preventing any collaboration from occurring. While TmeA-dependent activation of Arp2/3 via binding to N-WASP was demonstrated in *in vitro* pyrene-actin assays, evidence for *in vivo* relevance is lacking. However, an intriguing possibility is the involvement of TmeA in maintaining an active pool of N-WASP to activate Arp2/3 complexes bound along the length of actin filaments. However, this is speculative given that we did not observe N-WASP recruitment in prior live-cell imaging experiments that lasted for 360 s (Carabeo *et al*, 2007).

An added complexity is the potential involvement of growth factor receptors, e.g. EGFR, FGFR, integrins, etc., which could provide the signals to activate Arp2/3, and thus would be redundant to TarP and TmeA. Besides a potential compensatory role for growth factor receptors, an additional layer of complexity could come from signaling cross-talk. Given the plurality of factors involved in *Chlamydia* invasion, much more research is needed to identify their individual contributions and integrate them into a more comprehensive understanding of which signaling pathways influence collaboration between branched and filamentous actin nucleators.

Although TarP signaling has not yet been definitively linked to the activation of DRFs, we identified the recruitment of two DRFs, mDia1 and mDia2, which require activation by Rho-GTPases prior to serving as actin nucleators or filament elongation factors. Since TarP signals through Rac1 (Carabeo *et al*, 2007; Lane *et al*, 2008), which is known to activate mDia1 and mDia2 (51,52), our data support a model where TarP signaling contributes to the assembly of both branched and filamentous actin (Fig. 5B). Interestingly, Fmn1 recruitment was found to be TarP-dependent despite the fact that Fmn1 activity, unlike DRFs, is not contingent upon activation by Rho-GTPases. Fmn1 recruitment is likely mediated by the actin network nucleated by Arp2/3, with the barbed ends of each branch serving as binding sites, and thus may be indirectly recruited as a product of TarP-mediated actin remodeling. However, since Fmn1 recruitment was absent in TarP-deficient Chlamydiae, it seems that Arp2/3 which is activated by the alternate TmeA- dependent mechanism is incapable of compensating for the recruitment of Fmn1. Thus, the means by which Arp2/3 is activated appears to determine whether it will collaborate with Fmn1 during invasion. Nevertheless, the recruitment of multiple formin species highlights the importance of filamentous actin polymerization toward *Chlamydia* invasion, particularly since it is known that *Chlamydia* induces the assembly of hypertrophic microvilli, whose assembly is reliant upon the activity of formins (53–55). Finally, since the pan-formin inhibitor SMIFH2 is expected to inhibit all three formin species identified in our study (36), we are incapable of decoupling the effects of Fmn1, mDia1, and mDia2 in nucleator collaboration and subsequent actin remodeling. Overall, our data demonstrate the importance of formins toward *Chlamydia* invasion and introduce the possibility that TarP signaling contributes more to actin remodeling than was previously appreciated.

Throughout this study, we obtained evidence that actin dynamics associated with *Chlamydia* invasion have core and auxiliary elements. This is based on the correlation between the recruitment and turnover phases of actin dynamics. At the population level, actin dynamics, the effects of TarP-deficiency or inhibition of either nucleators led to the reduction in the rates of recruitment and turnover. However, upon detailed analysis where the recruitment phase for individual elementary bodies were matched with their corresponding turnover phase, we could not demonstrate significant correlation. Elementary bodies exhibiting rapid recruitment gave rise to rapid and slow turnover equally, and vice versa. When viewed from the perspective of an invasion process involving core and auxiliary components, the variability at the single-EB level could be accounted for by the accessory factors, which could include local concentration of actin, barbed ends, depolymerizing factors, actin crosslinkers, capping, etc., while an intact core component would explain the generally consistent trends observed at the population level. At the outset of invasion, *Chlamydia* makes contact and subsequently activates an assortment of host receptors (EGFR, PDGFR, ERα/β, β1-Integrin, etc.), which may influence the recruitment of actin, its nucleators, and actin depolymerization factors (Fig. 5A) (35,56,57). Furthermore, live-cell imaging of *Chlamydia* invasion did not reveal any consistent pattern of cell surface location of entry, instead occurring stochastically across the surface of the cell. As such, local variations in the levels of preassembled actin filaments or the concentration of signaling co-factors may exist. In addition, a myriad of other cellular machinery may be differentially accessible for exploitation by *Chlamydia*. What we observed, however, is that the loss of TarP had a substantial effect on the rate and distribution of actin kinetics, indicating that TarP is a core component that expands the repertoire of signaling events by coordinating the utilization of multiple accessory signaling proteins. Consequently, TarP coordinates the recruitment of a cohort of proteins which influence actin dynamics and contribute to efficient invasion of *C. trachomatis*, or vice versa where additional signaling events assist TarP in mediating efficient pathogen entry. In this context, TmeA, EGFR, PDGFR, ERα/β, β1-Integrin and others could be accessory signaling molecules that on their own are not sufficient to support efficient invasion and rapid actin dynamics. This is further evidenced by the fact that TarP deletion strongly attenuated pathogen internalization, indicating that activation of one or more of the above receptors and/or TmeA is insufficient to compensate for loss of TarP. Instead, their functional utility has to be in the context of TarP-dependent invasion.

## Materials and Methods

### Cell and Bacterial Culture

Green monkey kidney fibroblast-like Cos7 cells were cultured at 37 °C with 5% atmospheric CO2 in Dulbecco’s Modified Eagle Medium (DMEM; Gibco, Thermo Fisher Scientific, Waltham, MA, USA) supplemented with 10 µg/mL gentamicin, 2 mM L-glutamine, and 10% (v/v) filter-sterilized fetal bovine serum (FBS). Cos7 cells were cultured for a maximum of 15 passages for all experiments. McCoy B mouse fibroblasts (originally from Dr. Harlan Caldwell, NIH/NIAID) were cultured under comparable conditions. *Chlamydia trachomatis* serovar L2 (434/Bu) was propagated in McCoy cells and EBs were purified using a Gastrografin density gradient as described previously (58).

### Reagents

SMIFH2 (Tocris, Minneapolis, MN, USA) and CK666 (Sigma-Aldrich, St. Louis, MO, USA) were diluted upon receipt to 100mM stock concentration in DMSO, dispensed into single-use aliquots and stored at −20°C for no longer than 1 year after receipt. SMIFH2 was diluted to a working concentration of 10µM (1:10000) in supplemented DMEM, and CK666 was diluted to a working concentration of 100µM (1:1000) in supplemented DMEM.

### Invasion Assay

Efficiency of *C. trachomatis* invasion in Cos7 cells was performed as described previously (12). Briefly, Cos7 cells were seeded in 24-well plates containing acid-etched glass coverslips and allowed to adhere overnight. Cells were pretreated with SMIFH2 (10µM) (Tocris, Minneapolis, MN, USA), CK666 (100µM) (Sigma-Aldrich, St. Louis, MO, USA), or both, for 1 hour prior to infection. Following inhibitor treatment, cells were infected with EBs derived from wild-type *C. trachomatis* L2 (434/Bu), *C. trachomatis* in which TarP was deleted by FRAEM (ΔTarP), or *C. trachomatis* in which TarP expression was restored by *cis*-complementation (*cis* TarP) at MOI=50. EBs were allowed to attach to Cos7 cells for 30 min at 4°C. Cells were rinsed with cold HBSS before temperature shift to 37°C by the addition of pre-warmed DMEM + 10% FBS/2 mM L-glutamine and incubated at 37°C for 10 min. Cells were then washed with cold HBSS containing 100 μg/mL heparin to remove any transiently adherent EBs before fixation in 4% paraformaldehyde at room temperature for 15 min. Fixed cells were labeled with a mouse monoclonal anti-*Chlamydia* LPS antibody (BioRad CF 6J12, Hercules, CA, USA), rinsed with 1x PBS, and fixed once more in 4% paraformaldehyde for 10 min. Next, cells were permeabilized using 0.1% (w/v) Triton X-100 for 10 minutes at room temperature, rinsed with HBSS and labeled with rabbit polyclonal anti-*Chlamydia trachomatis* antibody (Abcam ab252762, Cambridge, MA, USA). Cells were then rinsed in 1x PBS and labeled with Alexa Fluor 488 anti-mouse (ThermoFisher #A28175, Waltham, MA, USA) and Alexa Fluor 594 anti-rabbit (ThermoFisher #A-11012) IgG secondary antibodies. Coverslips were mounted and observed on either a Leica DM500 (Leica, Allendale, NJ, USA) or Zeiss Axio Observer (Zeiss, Dublin, CA, USA) epifluorescence microscopes. Percent invasion efficiency was quantified as total EBs (red) – extracellular EBs (green)/total EBs (red) x 100%.

### Quantitative live cell imaging of *Chlamydia* invasion

Cos7 cells were seeded onto Ibidi µ-Slide 8-well glass-bottomed chambers (Ibidi, Fitchburg, WI, USA) and allowed to adhere overnight prior to transfection. Cells were transfected with fluorescent proteins as indicated, using Lipofectamine 3000 (Thermo Fisher, Waltham, MA, USA) according to manufacturer directions. Transfection was allowed to proceed overnight before replacing media with fresh DMEM + 10% FBS/2 mM L-glutamine and allowing protein expression to continue for a total of 24 hours post-transfection. Transfection was verified on either a Leica SD6000 or Nikon CSU-W1 (Nikon, Melville, NY, USA) spinning disk confocal microscope prior to application of DMEM containing SMIFH2 (10µM), CK666 (100µM) or both. Wells were individually infected with CMTPX-labeled wild-type *C. trachomatis* L2 (434/Bu), unless otherwise indicated, at MOI=20 and promptly imaged using a 60x objective (NA 1.40) in a heated and humidified enclosure. Images were collected once every 20 seconds for 30 minutes, with focal plane maintained using an infrared auto-focusing system. Upon completion of the imaging protocol, the next well was infected and imaging repeated; mock-treated wells were imaged first to allow inhibitor treatment sufficient time to achieve inhibition. Images were compiled into videos using NIH ImageJ and analyzed to identify protein recruitment events. The mean fluorescence intensity (MFI) of such events was measured for each timepoint alongside the local background MFI of a concentric region immediately outside the recruitment event. Background MFI was subtracted from recruitment MFI for each timepoint and converted into fold recruitment by dividing the MFI of each timepoint against the baseline MFI, defined as the average MFI of five timepoints (100 sec) prior to the start of recruitment.

### Plasmids and DNA preparation

pEGFP-Actin–C1 (59) was provided by Dr Scott Grieshaber (University of Idaho). pEGFP-N1-ACTR3 (Arp3- GFP) (38) was obtained from Dr Matthew Welch (Addgene plasmid #8462). Arp3-pmCherryC1 (60) was a gift from Christien Merrifield (Addgene plasmid #27682). mEmerald-mDia1-N-14 (Addgene plasmid#54157), mEmerald-mDia2-N-14 (Addgene plasmid #54159), and mRuby-LifeAct-7 (Addgene plasmid#54560) were gifted by Michael Davidson. pEGFPC2-FmnIso1b (61) was a gift from Philip Leder (Addgene plasmid #19320). pmCherryC1-FmnIso1b was generated by a variant of Gibson assembly termed FastCloning (62) and included removal of EGFP from pEGFPC2-FmnIso1b and insertion of mCherry derived from Arp3-pmCherryC1. Kanamycin-resistant transformants were selected and propagated for plasmid isolation prior to sequence verification using an mCherry-Fwd sequencing primer supplied by Eurofins genomics. All plasmids were isolated using MiniPrep DNA isolation kits (Qiagen, Valencia, CA, USA) following a variant protocol for DNA isolation termed MiraPrep (63). Following plasmid isolation, the eluate was precipitated by addition of 3M sodium acetate (Invitrogen, Waltham, MA, USA) at 10% (v/v) of eluate volume followed by addition of 250% (v/v) absolute ethanol calculated after addition of sodium acetate. The mixture was incubated at 4°C overnight and centrifuged at 14,000×g for 15 minutes at 4°C. Supernatant was removed and 70% ethanol was added, followed by centrifugation at 14,000×g for 10 minutes at 4°C. Supernatant was removed once more, and precipitated DNA was resuspended in nuclease-free H_2_O. Primers for assembly of pmCherryC1-FmnIso1b are as follows: (pEGFPC2) Fwd: 5’ GGCGGAAGCGGAAGC<u>ATGGAAGGCACTCACTGCACC</u>3’, Rev: 5’GCCCTTGCTCACCAT <u>GGTGGCGACCGGTAGCG</u> 3’; (pmCherryC1) Fwd: 5’ CTACCGGTCGCCACC<u>ATGGTGAGCAAGGGCGAGG</u> 3’, Rev: 5’ GTGAGTGCCTTCCAT<u>GCTTCCGCTTCCGCCG</u> 3’.

### Graphs and statistical analysis

Violin plots were made using the ggplot2 base package (version 3.1.0) as a component of the Tidyverse package (https://cran.r-project.org/web/packages/tidyverse/index.html) in rStudio (version 4.0.3). Kolmogorov-Smirnov tests to determine statistical significance between violin plots was performed using the dgof package (version 1.2, https://cran.r-project.org/web/packages/dgof/index.html) in rStudio. Tabulated statistical analyses of violin plots were generated using base R statistics and the moments package (version 0.14, https://cran.r-project.org/web/packages/moments/index.html) in rStudio and assembled in Excel (Microsoft, Redmond, WA, USA). Linear regression analyses, recruitment plots, invasion assays, and all statistics associated with these data (pairwise T-test, correlation coefficient, SEM) were performed in Excel. All graphs were assembled using the free and open-source software GNU Image Manipulation Program (GIMP, https://www.gimp.org/) and Inkscape (https://inkscape.org/). Proposed model for collaboration (Fig. 5) was assembled using BioRender (https://app.biorender.com/).

## Acknowledgements

We thank the members of the Carabeo lab for extensive feedback regarding the design, direction, and analysis of the study. We thank Kenneth Fields (University of Kentucky College of Medicine) for the kind gift of both ΔTarP and TarP *cis*-complemented strains. This study was supported by funding from the U.S. National Institutes of Health, National Institutes of Allergy and Infectious Disease grants R01 AI065545 (R.A.C.). M.D.R. was further supported by a fellowship from the Seattle Chapter of Achievement Rewards for College Scientists (ARCS). The contents and views expressed within this publication are the sole responsibility of the authors.

## Author Contributions

M.D.R. and R.A.C. designed the experiments and wrote the manuscript. M.D.R. performed the experiments, and M.D.R. and R.A.C. analyzed the data.

## Data Availability

Datasets regarding the analysis of invasion assays, fluorescence microscopy and associated kinetic analyses are available from the Dryad repository https://doi.org/10.5061/dryad.0k6djhb0g.

